# Modelling sex-specific crossover patterning in Arabidopsis

**DOI:** 10.1101/440255

**Authors:** Andrew Lloyd, Eric Jenczewski

**Affiliations:** Institute of Biological, Environmental and Rural Sciences, Aberystwyth University, Aberystwyth, SY23 3EB, UK; Institut Jean-Pierre Bourgin, INRA, AgroParisTech, CNRS, Université Paris-Saclay, 78000, Versailles, France; INRA UMR1349 Institut de Génétique, Environnement et Protection des Plantes, France

**Keywords:** recombination, crossovers, interference, beam-film, sex-specific

## Abstract

Interference is a major force governing the patterning of meiotic crossovers. A leading model describing how interference influences crossover-patterning is the beam film model, a mechanical model based on the accumulation and redistribution of crossover-promoting stress along the chromosome axis. We use the beam-film model in conjunction with a large Arabidopsis reciprocal back-cross data set to gain mechanistic insights into the differences between male and female meiosis and crossover patterning. Beam-film modelling suggests that the underlying mechanics of crossover patterning and interference are identical in the two sexes, with the large difference in recombination rates and distributions able to be entirely explained by the shorter chromosome axes in females. The modelling supports previous indications that fewer crossovers occur via the class II pathway in female meiosis and that this could be explained by reduced DNA double strand breaks in female meiosis, paralleling the observed reduction in synaptonemal complex length between the two sexes. We also demonstrate that changes in the strength of suppression of neighboring class I crossovers can have opposite effects on effective interference depending on the distance between two genetic intervals.

## INTRODUCTION

Meiotic crossovers shuffle parental genetic information generating new combinations of alleles. In most species the presence of one crossover inhibits nearby crossover formation so that inter-crossover distances are greater and more uniform, than if placed at random. This phenomenon, crossover interference, was first noted in genetic studies over a century ago (1, 2), however it is only in the last few years that insights into its mechanistic basis have begun to surface.

Studies in Arabidopsis (3), yeast (4) and humans have shown that the inhibitory effect of interference spreads across a defined physical chromosome distance. This is usually measured in μm synaptonemal complex (SC) although there is some evidence that it is mediated by the chromosome axes (whose lengths are proportional to that of the SC) prior to synapsis (reviewed 5). In yeast, interference is, at least in part, mediated by Topoisomerase II (4) and wt levels of interference require SUMOylation and subsequent ubiquitin-mediated removal of TopoII and also of the axis component Red1/Asy3 (4). These findings are consistent with suggested roles for the chromosome axis and local stress relief via DNA remodeling, in mediating interference.

Several approaches have been used to model CO patterning, the most notable being the gamma model and the beam-film model. The gamma model is a statistical model based on the observation that the distances between two crossovers are relatively uniform, following a gamma distribution. Under this model “effective interference strength” is highest when distances between crossovers show the least variation. This results in a large value of the gamma shape parameter.

In contrast, the beam-film model is a mechanistic model whose various parameters have biological correlates. In the beam-film model, each bivalent has a number of precursor sites (DSBs) that are subject to mechanical stress. CO maturation at precursor sites is promoted by stress and this stress is relieved locally following CO-designation. As stress promotes COs, stress relief propagating out from crossover sites inhibits the formation of additional COs nearby. In the beam-film model, interference strength is highest when stress relief propagates furthest from designated crossover sites.

In most species, there are multiple crossover pathways. The majority of crossovers occur via the interference sensitive class I pathway and are dependent on the ZMM group of proteins identified initially in yeast (Zip1, Zip2, Zip3, Zip4, Mer3, Msh4, and Msh5) (6). Crossovers occurring via this pathway are specifically marked by Zip3/Hei10 and MLH1 foci at late pachytene (7–9). A number of secondary “clean-up” pathways repair DSBs not metabolized by the class I pathway (10, 11). These clean-up pathways mostly repair DSBs as noncrossovers, but also contribute a smaller number of crossovers (i.e. class II crossovers) which are insensitive to interference (12) and usually make up 10-15% of total recombination events (e.g. 7, 12, 13).

In their simplest forms, the gamma and beam-film models of crossover patterning deal exclusively with class I crossovers and therefore experimental data used in comparisons with simulated data must be class I specific, e.g. Zip3/Hei10 foci distributions. Crossover patterning in yeast has been extensively explored using the single-pathway beam-film model and the effects of different parameters on crossover distributions are now well understood. Consideration of class I crossovers alone, however, is not always possible or appropriate. For example, cytological analyses of female meiosis in Arabidopsis remain challenging and, as the number of crossovers per chromosome is low, well over a thousand cells would need to be analyzed to achieve the same number of inter-interval distances (the limiting factor for analyses) commonly reported for yeast chromosomes (4, 14). Cytological data are also not appropriate when wanting to consider patterning of both class I and class II crossovers, as there is as yet no robust cytological marker for class II crossovers. In these instances, beam-film best-fit simulations must be used in combination with genetic data, which include the result of both class I and class II crossovers. Backcross populations, for example, are particularly well suited to this kind of analysis.

Using a large Arabidopsis reciprocal backcross recombination data set (~1500 individuals, ~380 markers for both male and female) it has been shown using the two-pathway gamma model that crossover interference is higher in female meiosis than in male meiosis, that male meiosis has a higher proportion of non-interfering COs and that class-II COs are non-uniformly distributed (15).

We use the two-pathway beam-film model to analyze the same large Arabidopsis dataset, identifying parameter best fits to gain mechanistic insights into the differences between male and female crossover patterning. Using the best-fit parameter values identified as a starting point we then model the influence of various parameters on crossover patterning under the two-pathway beam-film model.

## MATERIALS AND METHODS

### Experimental data

Experimental dataset used has been previously published (16) and was derived from large Arabidopsis reciprocal backcross populations. On average, 1,505 plants were genotyped for 380 SNPs in the male population and 1,507 plants genotyped for 386 SNPs in the female population (380 in common). As the average distance between markers is small in this data set – 316 kb in male, 311 kb in female – the number of double crossovers (DCOs) in a single interval are expected to be negligible. It was therefore assumed during analysis that all recombination events were identifiable.

### Beam film parameter optimization

Beam-film simulations were performed and best-fit parameters determined using MADpatterns (17) and custom perl scripts with an approach based on that described in (14). For each chromosome and each sex at least three rounds of analysis were undertaken. In each round of each analysis 30,000 bivalents were simulated for a range of parameter values. In the first round, relatively broad value ranges of optimised parameters (*Smax:* 2 – 10 L: 0.4 – 1.7; *T2Prob*: 0.002 – 0.008; *cL*: 0.3 – 1.3 and *cR*: 0.3 – 1.3) were chosen based on values described in Zhang et al (2014) and comparison of ad hoc simulations with analysis of experimental datasets (15). Parameters *N, B, E, Bs/Be/Bd, A* and *M* were set at appropriate default values (see below). In the next two rounds, progressively smaller step-sizes between values were used to arrive at the final parameter values.

For each round of analysis, the crossover distributions, CoC curves and event distributions (distribution of number of COs per gamete) simulated for each chromosome were determined using MADpatterns (17) and compared to those obtained for the relevant sex and chromosome from the experimental data set; number of intervals = 13. Importantly the experimental data are gamete data, while the MADpatterns program simulates (and outputs) bivalent data (i.e. all crossovers on a pair of homologous chromosomes). Therefore, all simulated bivalent crossover frequencies were halved to convert to gamete crossover frequencies. Bivalent event distributions were also converted to gamete event distributions, assuming random assignment of each crossover to two of the four chromatids. Parameter sets were ranked based on the difference between simulated and experimentally determined CoC distributions [Scores_CoC_ = Σ_IID_ abs(log2(CoC_sim/_CoC_exp_))], CO distributions [Scores_CO_ = Σ_int_ (CO_sim_ − CO_exp_)^2^] and event distributions [Score_ED_ = Σ_Enum_ (ED_sim_ − ED_exp_)^2^]. Simulations were ranked for each score and final parameter values chosen were those with the lowest rank-sum.

### Fixed Parameter Values

Beam-film model parameters *N, B, E, Bs/Be/Bd, A* and *M* were fixed based on known values of the biological correlates, parameters that tend to be stable between species (14), or suggested default values (18). A description of each of these parameters and values used is given below, further explanations can be found in refs. (14, 17).

#### N – Precursor sites per bivalent

Each meiotic double strand break (DSB) corresponds to a recombination precursor site. In Arabidopsis the total number of DSBs per meiosis is estimated at around 250 (19). For beam-film modelling, the number of DSBs for each chromosome was calculated as 250 multiplied by the proportion of the total genome size (in Mb) contributed by that chromosome (Table S1.). To simulate reduced DSB formation in female meiosis, total DSB number was set at 150 and the per-chromosome number of precursors calculated as above.

#### B – Similarity in total precursor number between bivalents

*B* sets the similarity of precursor number between the multiple bivalents simulated for a given chromosome in each round of analysis: 0 = Poisson, 1 = constant. When *B* ≠ 1, i.e. precursor number is not constant, parameter *N* specifies the average number of precursors per bivalent. *B* was set to 1 for all simulations.

#### E – Evenness of precursor spacing

*E* sets the evenness of precursor spacing: 0 = random, 1 = even. There is considerable experimental evidence that DSB spacing is non-random (20, 21). For numerous organisms a parameter value of 0.6 has been found appropriate (14). *E* was set to 0.6 for all simulations.

#### Bs/Be/Bd – Recombination “black hole” start/end/precursor density

Recombination black hole start and end points correspond to the start and end of the recombination suppressed centromeric region. These values were determined based on the recombination frequencies observed in the backcross data (Table S1, Figure S1) and correspond to regions with high DNA methylation and low H3K4me3 indicative of heterochromatin (21).

#### Bsmax – Similarity in maximum stress levels between bivalents

*Bsmax* sets the similarity of *Smax* between simulated bivalents: 0 = Poisson, 1 = constant. When *Bsmax* ≠ 1, i.e. maximum stress levels are not constant, parameter *Smax* specifies the average maximum stress per bivalent. *Bsmax* was set to 1 for all simulations.

#### A – Intrinsic precursor sensitivities

Parameter *A* determines how precursor sensitivities are assigned. For all simulations *A* was set to 1 – sensitivities assigned from a uniform distribution.

#### M – Crossover maturation efficiency

In the beam film model, it is possible to model failure of crossover maturation. If failure occurs, the CO-designated site inhibits nearby crossovers but does not itself develop into a crossover. 0 = no maturation, 1 = 100% maturation. *M* set to 1 for all simulations.

### Double crossover class determination

The proportion of each class of DCO for a given inter-interval distance was determined from simulations modelling the formation of class I crossovers only (*T2Prob* = 0), class II crossovers only (*Smax* = 0), or both class I and II crossovers. For each simulation, numbers of DCOs were tallied for each inter-interval distance (IID, the distance between a pair of genetic intervals). For each IID, numbers of DCOs involving two class I COs (DCO_I_I_), two class II COs (DCO_II_II_) or all DCOs (DCO_ALL_) were calculated from the respective simulations. DCO_I_II_ = DCO_ALL_ − (DCOI_I + DCO_II_II_).

### Response of model to parameters *L, Smax, T2Prob* and N

To investigate the response of the model to parameters *L*, *Smax* and *T2Prob* we simulated 30000 bivalents for an “idealized” male Arabidopsis chromosome (*N* = 60, *B* = 1, *E* = 0.6, *Bs* = 0.45, *Be* = 0.55, *Bd* = 0.01, *Smax* = 9, *Bsmax* = 1, *A* = 1, *L* = 0.7, *cL* = 0.8, *cR* = 0.8, *M* = 1, *T2Prob* = 0.004) as described above, varying one specified parameter.

## RESULTS

### Beam-Film Parameter Best-Fit Simulations

We used the MADpatterns program (18) and custom scripts to simulate crossover patterning and identify parameter best-fits for each of the five Arabidopsis chromosomes for both male and female meioses. Parameters *N, B, E, Bs/Be/Bd Bsmax, A* and *M* were fixed for a given set of simulations (see methods), while *Smax, L, T2Prob* and *cL/cR* were explicitly optimized. Best-fit values were identified by comparing crossover number and distribution, and interference relationships (CoC curves) of simulated and experimental datasets (Figure 1 & S1-2). Simulations using best-fit parameters gave good approximations of the experimental data (Figure 1 & S1-2).

**Figure 1.**
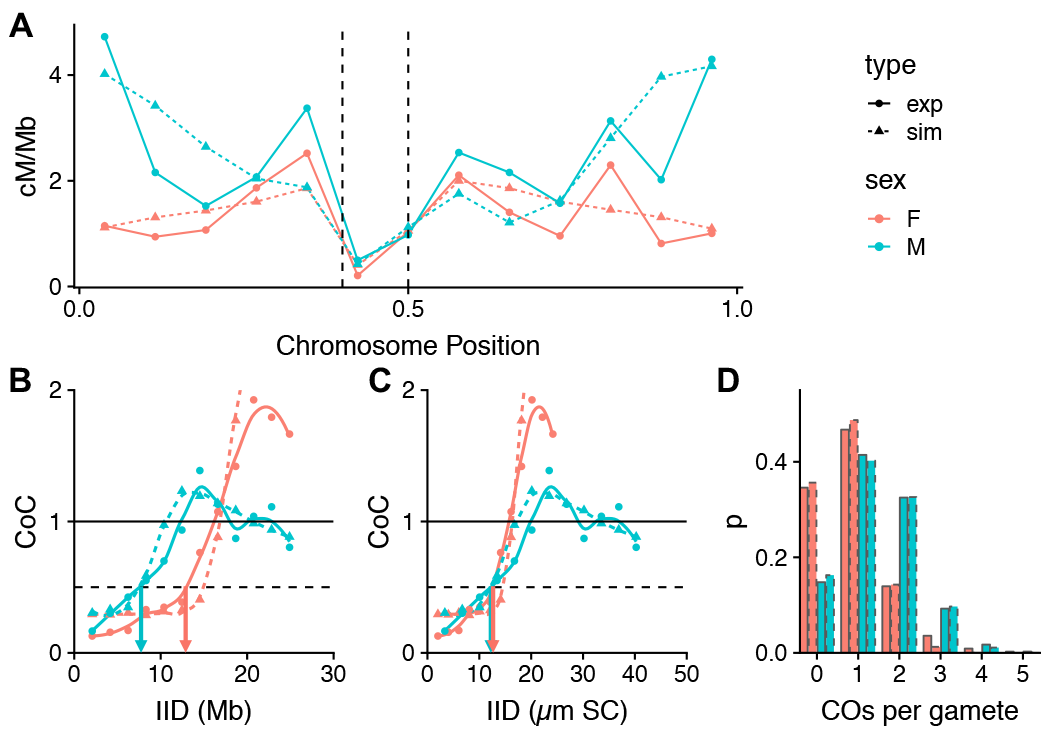
Crossover analysis for Arabidopsis chromosome 5. Each analysis includes experimental (solid lines) and simulated (dashed lines) data from male (blue) and female (orange) meioses. **A** Crossover distributions for Arabidopsis chromosome 5. Dashed lines represent the limits of the centromeric region over which precursor number is markedly reduced during simulations. **B-C** CoC curves for chromosome 5 with inter-interval distance (IID, the distance between a pair of genetic intervals) measured in Mb (**B**) or μm SC (**C**). L_CoC_ positions for male and female are indicated by blue and orange arrows respectively. **D** Event distribution for chromosome 5.

### Optimized Parameters

#### Smax - Maximum stress level per bivalent

In the beam-film model, whether or not a precursor site (DSB) develops into a crossover is determined by the crossover-promoting “stress” (*S*) experienced by that precursor as well as the precursor’s sensitivity (determined by parameter *A*, see methods). The biological correlate of *Smax* i.e. the nature of the crossover promoting stress is not precisely defined but may relate to the expansion of chromatin during early prophase (22). When simulating each bivalent, the value of *S* is progressively increased until *S = Smax.* At some point *S* reaches a critical value at which the most sensitive precursor undergoes CO-designation and stress-relief extends out from that position. As *S* increases to *Smax*, additional precursors usually experience sufficient stress to promote the designation of further COs. At any point during the simulation, the actual stress experienced by a precursor is the current value of *S*, minus the sum of any stress-relief due to interference from nearby COs. Optimum values of *Smax* for the five chromosomes were similar in males and females: male 7 ± 1.9 and female 6.9 ± 0.7, p = 1 (Bonf. corrected) (Figure 2, Table S1).

#### L_BF_ – Stress relief distance

The parameter *L*_*BF*_ corresponds to the length of the chromosomal interval over which a CO relieves stress i.e. stress-relief propagates out from COs a distance of ½ *L*_*BF*_ in either direction. The magnitude of the stress-relief decreases exponentially with distance from the CO, such that there is maximal stress-relief in the middle of the interval (i.e. immediately surrounding the CO) and almost no stress-relief at either end of the interval (14, 22).

When running simulations *L*_*BF*_ is specified as the proportion of total chromosome length (i.e. chromosome length is set to 1), but is converted to length in Mb or μm SC to enable comparisons between chromosomes of different lengths. When measured in Mb (*L*_*BF_Mb*_) best-fit estimates of *L*_*BF*_ were significantly higher in females: *L*_*BF_Mb*_ − male 17.1 ± 3.5 Mb and female 28.8 ± 3.1 Mb, p = 0.0095 Bonf. corr. (Figure 2, Table S1). When the distance metric was converted to μm SC (*L*_*BF_SC)*_), using the best available estimates of SC length in the two sexes (3), there was no-longer any difference in *L*_*BF*_ between the two sexes: *L*_*BF_CoC*_ − male 27.7 ± 5.6 μm and female 23.7 ± 2.5 μm, p = 1 Bonf. corr. (Figure 2). For some chromosomes, optimum values of *L*_*BF*_ were larger than the chromosome in question. This is expected if one CO suppresses the formation of additional COs more than half the length of the chromosome away. An example can be seen for chromosome 2 in females which has an estimated SC length of 16.2 μm and an *L*_*BF_CoC*_ estimate of 25.9 μm. As can be seen from the CoC curve for this chromosome (Figure S2) it is clear that the observed number of DCOs are less than expected (i.e. CoC < 1) even when intervals are at opposite ends of the chromosome (e.g. inter-interval distance ~14 μm).

#### cL/R – Left and Right end clamping

In the beam-film model, “clamping” at chromosome ends determines how stress is supported in terminal regions. Unclamped chromosome ends (cL = 0; cR = 0) cannot support stress and so locally relieve stress, behaving as if there were a crossover at the chromosome end. Clamped chromosome ends (cL = 1; cR = 1) experience stress as elsewhere along the bivalent. As the interference signal cannot come from beyond the end of the chromosome, recombination frequencies will tend to be higher at the end of chromosomes than for internal regions when clamped. Total clamping averages (cL/R) for male and female were calculated from the estimated values of cL and cR for each sex. Clamping values were variable between chromosomes but there was no significant difference between the average clamping values, male 0.78 ± 0.16 and female 0.69 ±0.13, p = 1 Bonf. corr. (Figure 2).

#### T2Prob - Probability that a non-crossover designated precursor will form a Type II crossover

The average optimized T2Prob values were significantly higher in male than female meiosis, 0.0063 ± 0.0010 and 0.0036 ± 0.0008 respectively (p = 0.026, Bonf. corrected, Table 1, Figure 2). This value corresponds to the proportion of non-crossover designated precursor sites that become Type II COs and assumes equal numbers of precursor sites i.e. meiotic DSBs in male and female meiosis. We also determined what proportion of the total number of crossovers occur via the class II pathway (i.e. p = CO_II_/ (CO_I_+ CO_II_). These values were equivalent for the two sexes: 0.14 ± 0.02 male, 0.14 ± 0.03 female, p = 1 Bonf. corr.

**Figure 2.**
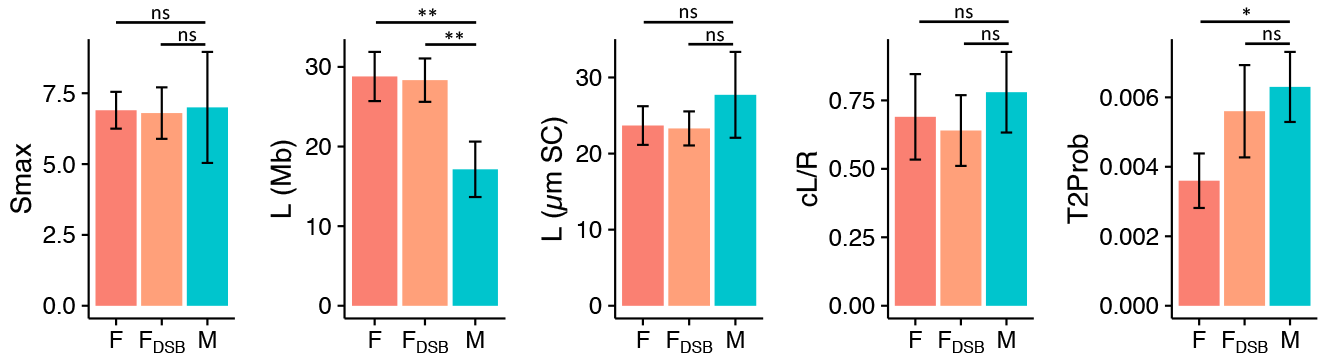
Beam-film best-fit parameter values. Standard simulations for male (M) and female (M) assumed 250 DSBs per meiosis. Simulations for female were run a second time assuming 150 DSBs (F_DSB_). * p < 0.05, ** p < 0.01, after Bonferroni multiple comparison correction.

### Beam-Film Parameter Best-Fit Simulations – Modelling reduced DSB formation in females

While there are relatively good estimates for the number of DSBs in male meiosis in Arabidopsis, cytological analyses of female meiosis are more challenging and there are no reliable estimates of DSB numbers. Thus, while we have assumed equal numbers of DSBs in male and female meiosis, it is possible that DSB numbers differ between the two sexes. Meiotic DSBs occur in loop DNA that has been recruited to the chromosome axis (23). In Arabidopsis female meiosis there are fewer (albeit larger) chromatin loops and the chromosome axis is 40% smaller than in male meiosis (3) which could feasibly result in a similar reduction in DSBs (24, 25). To understand whether reduced DSB numbers would have any effect on crossover patterning and/or estimates of parameter values in female meiosis, we repeated the best fit simulations assuming a reduction in DSBs equal to the reduction in SC length i.e. approx. 40% reduction, or 150 (rather than 250) DSBs per meiosis.

Optimised values of *Smax*, *L*_*BF_Mb*_, *L*_*BF_SC*_ and cL/cR were identical for both sets of simulations (F and F_DSB_, Figure 2). Estimates of *T2Prob* were higher for simulations of female meiosis with reduced DSB numbers, and the optimized value no-longer differed from the value of *T2Prob* estimated for male meiosis (Figure 2). In the beam film model, the parameter *T2Prob* is the number of non-crossover designated DSBs that become class II COs. Thus, despite the difference in the parameter value estimates of *T2Prob* for female meiosis with 250 or 150 DSBs, the total number of class II COs estimated using both approaches is the same: 250 DSBs × *T2Prob* 0.0036 ± 0.0006 = 0.87 ± 0.15 COs; 150 DSBs × *T2Prob* 0.0056 ± 0.0010 = 0.90 ± 0.17 COs.

### Crossover Distribution Analysis

Simulated CO distributions closely matched those observed in the experimental data with highest recombination in males in distal regions and highest recombination in females closer to the centromere. (Figure 1, Figure S1). The exception was the short arms of chromosomes 2 and 4 in males which have high experimental recombination rates but had low levels of recombination when simulated using the global best-fit parameters (Figure S1. Sim vs Exp – COfreq all Chrs). It is possible that this is related to the presence of the NORs on the short arms of chromosomes of these two chromosomes.

### CoC Analysis

The coefficient of coincidence (CoC) is the ratio of the observed and expected numbers of double crossovers (DCOs) for a given pair of intervals, given the rates of single COs in the two intervals. When interference strength is high, CoC values tend to be low as there are fewer DCOs observed than expected (one CO suppresses the occurrence of nearby COs). CoC shows a characteristic curve when plotted against inter-interval distance (Figure 1B-C), with low CoC for small inter-interval distances and CoC approximating 1 for large inter-interval distances. A useful measure when analyzing such curves is *LCoC*, the inter-interval distance at which the curve crosses the line CoC = 0.5 (dashed line, Figure 1B-C). For all analyses the simulated data gave *L*_*CoC*_ values that were no different from those determined from the equivalent experimental data (Table 1.). For both experimental and simulated data, *L*_*COC*_ was significantly smaller in males than in females if measured in Mb but showed no difference between when measured in μm SC (Table 1, Figure 1 and Figure S2).

**Table 1.** *L_CoC_*values

| | Mb | | | $\mu m$ SC | | |
| --- | --- | --- | --- | --- | --- | --- |
|  | male | female | p value<sup>#</sup> | male | female | p value<sup>#</sup> |
| experimental | 7.05 ± 0.50 | 12.84 ± 1.50 | 7.90E-07 | 11.65 ± 0.86 | 12.83 ± 1.50 | 1 |
| simulated | 6.30 ± 1.05 | 11.60 ± 0.83 | 1.40E-05 | 10.21 ± 1.75 | 11.20 ± 0.78 | 1 |
| p value<sup>#</sup> | 1 | 1 |  | 1 | 1 |  |

<sup>#</sup>Bonferroni corrected

### Response of the two-pathway beam-film model to parameters L, *Smax* and *T2Prob*

To further investigate how changes in individual parameters affect the two-pathway beam-film model we simulated crossovers using a range of parameter values.

#### crossover Distributions

As the distance over which crossover interference is propagated (*L*_*BF*_) decreased, crossover frequencies increased. The change in frequency was particularly marked in distal regions although all regions experienced some increase in crossovers (Figure 3A). When *L*_*BF*_ was reduced below half of chromosome length, increases in crossovers was also strong adjacent to the centromere (Figure 3A).

Changes in *Smax* – the maximum crossover promoting force (or “stress”) experienced by recombination pre-cursors – had similar effects on crossover distribution as those observed for changes in *L*_*BF*_. Higher values of *Smax* (like low values of *L*_*BF*_) resulted in large increases in recombination in distal regions, but recombination rates were largely unchanged in proximal regions (Figure 3D). For low *Smax*, crossover rates were very low near the ends of the chromosomes and resulted in the most proximal crossover distribution (Figure 3D). Increases in *T2Prob* resulted in a small uniform increase in COs across the chromosome (Figure 3G).

#### CoC

Increases in *L*_*BF*_ shifted the CoC curve to the right resulting in increased *L*_*COC*_ values (Figure 3B). For small inter-interval distances increases in *L*_*BF*_ resulted in no change or a small increase in CoC (Figure 3C).

For low values of *Smax*, CoC curves were shifted more to the right and for higher values of *Smax* CoC curves were shifted to the left, however the effect on *L*_*CoC*_ was not as pronounced as observed for changes in *L*_*BF*_ (Figure 3B & 3E). Again, for small inter-interval distances decreases in *Smax* resulted in no change or an increase in CoC (Figure 3F). As has been observed previously in yeast (14), *Smax* had less influence on CoC than L.

Changes in *T2Prob* had limited effect on CoC for intermediate and large inter-interval distances, with no broad effects on CoC curves and no change in *L*_*CoC*_ values (Figure 3H). For small inter-interval distances, however, the level of class II crossovers had a marked effect, with higher number of class II crossovers resulting in higher values of CoC (Figure 3I).

Thus for smaller inter-interval distances the main determinant of CoC is the number of class II crossovers, while for intermediate and large inter-interval distances CoC values are mainly determined by *L*_*BF*_ and to a lesser extent *Smax* i.e. factors affecting (interference sensitive) class
I crossovers.

**Figure 3.**
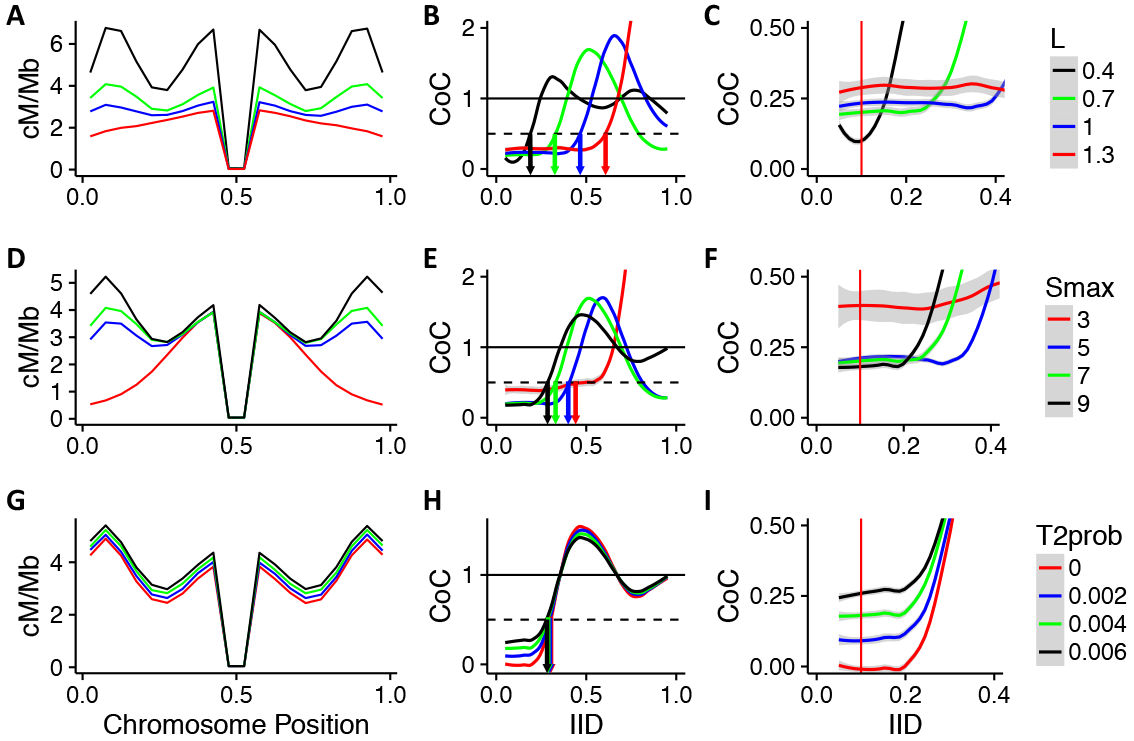
Effect of beam-film parameters on crossover patterning in Arabidopsis. The effect of altering a single beam film parameter – L (**A-C**), Smax (**D-F**) or T2Prob (**G-I**) – on crossover distribution (**A, D & G**) and CoC (**B-C**, **E-F** and **H-I**). Red vertical lines in **C, F** and **I** represent IID = 0.1.

### Differing classes of double COs at small and large IIDs cause opposing effects of interference on CoC

One of the less intuitive outputs of the analysis above was that for small inter-interval distances, an increase in the distance over which the interference signal is propagated can result in increased values of CoC (Figure 3C). To further understand this phenomenon, we sought to identify how changes in *L*_*BF*_ might differentially affect the expected and observed number of double COs (the determinants of CoC) for different IIDs. Beam film simulations demonstrated that increased *L*_*BF*_ resulted in a small decrease in the expected number of double COs (DCOs) for small and large IIDs (IID = 0.1 and 0.5; Figure 4A). This is unsurprising given that the expected number of DCOs for a pair of intervals is purely based on the respective rates of COs in the two intervals. In contrast, the observed number of DCOs changed dramatically for IID = 0.5, but only marginally for IID = 0.1 (Figure 4A) in response to changes in *L*_*BF*_. As a result, CoC dramatically decreased for IID = 0.5 with increased *L*_*BF*_ but increased slightly for IID = 0.1 (Figure 4B).

We reasoned that the difference in behavior might be due to the nature of the DCOs formed at smaller and larger IIDs. For example, DCOs can occur between two class I COs, two class II COs or between a class I and a class II CO but interference only directly suppresses those involving two class I COs. We therefore ran beam film simulations with class I COs only (T2Prob = 0), class II COs only (Smax = 0), or both class I and class II COs and determined numbers of the different classes of DCOs formed for each set of simulations (Figure 4C). From these numbers we determined the proportions of the different classes of DCOs (Figure 4D) that occur for different IIDs under standard conditions (i.e. when simulating both class I and class II COs). For small IIDs DCOs are almost exclusively formed between a class I CO and a class II CO (Figure 4D). In contrast, for larger IIDs (≥ 0.4) the majority of DCOs are formed between two class I COs (Figure 4D). Cytological observations in tomato reporting the same phenomenon (26) suggest this is a general feature of meiosis. As interference only suppresses DCOs involving two class I COs, changes in *L*_*BF*_ will only directly affect DCO formation at larger IIDs. This pattern holds when the proportion of class II crossovers falls within the range normally observed (5-20%), although when class II crossovers are absent or make up the majority of crossovers then most DCOs involve two class I or two class ll COs respectively for all IIDs (Figure S3).

Both the expected number of DCOs and observed DCOs at small IIDs are indirectly affected by increased *L*_*BF*_ due to the associated decrease in the frequency of class I COs. The magnitude of the change is greater for the expected number of DCOs, which can be seen from the equations below. Here *CI* and *CII* are the rates of class I and class II crossovers respectively in the two intervals:

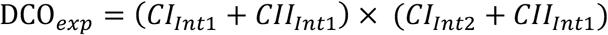

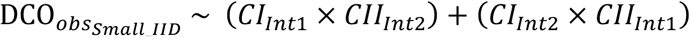

For small IIDs, while *CI* ≫ *CII*, the reduction in the expected number of DCOs is approximately twice that of the observed reduction in DCOs, resulting in an increase in CoC.

**Figure 4.**
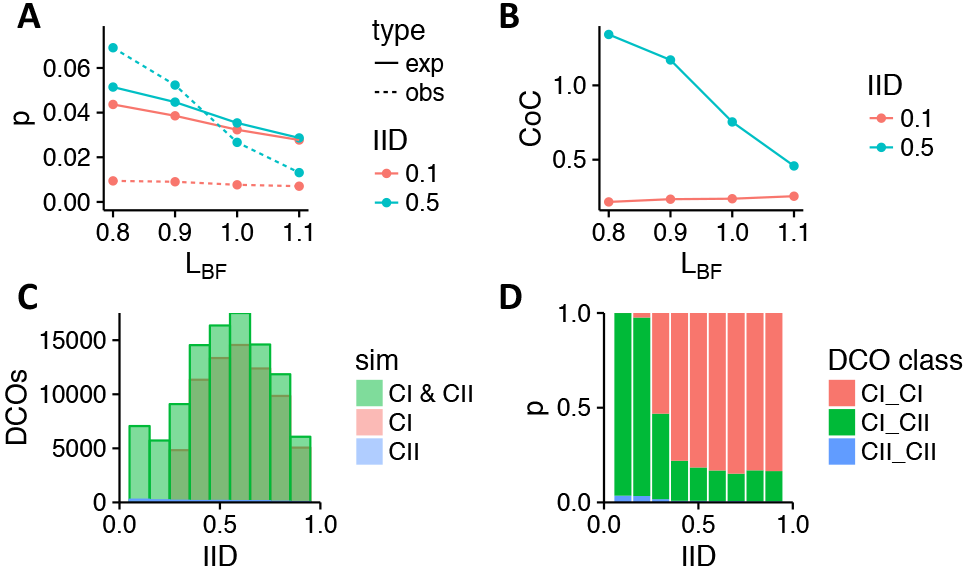
Influence of IID on CoC response to changes in *L*_*BF*_. **A** The expected (solid line) and observed (dashed line) proportion of interval pairs receiving a double crossover (DCO) for two different inter-interval distances (IIDs); calculated from simulations with varying values of *L*_*BF*_. **B** CoC values for two IIDs calculated from simulations with varying values of *L*_*BF*_. **C** The number of DCOs observed for different IIDs from simulations involving class l and class ll crossovers (Cl & Cll), class l crossovers only (CI) or class ll crossovers only (CII). **D** The proportions of DCOs formed between two class I crossovers (CI_CI), two class II crossovers (CII_CII), or a class I and a class II CO (CI_CII) for different IIDs.

### Response of the two-pathway beam-film model to pre-cursor (DSB) numbers

As described above a 40% decrease in DSB numbers had few effects on estimates of beam-film parameters. To further investigate the effect of DSBs on crossover patterning in Arabidopsis we simulated crossovers for a range of DSB numbers. As well as simulating wt meiosis, we also modelled the effect of altered DSB numbers in situations where a large proportion of DSBs become class II crossovers, such as is observed in some mutant contexts (27).

Altered DSB number had relatively little effect on crossover distributions or CoC curves in a wt context (Figure 5A-B). For CO distribution, the main change was a general flattening of the distribution with lower DSB number. The only clear difference in CoC was for small inter-interval distances, where higher DSB numbers resulted in higher values of CoC (Figure 5C). In contrast, altering the number of DSBs in a context where a high proportion become class II crossovers had a dramatic effect on crossover patterning. Here increased DSBs resulted in proportionate increases in crossovers (Figure 5D). Regardless of the number of DSBs, CoC values were approximately 1 for all inter-interval distances (Figure 5E-F).

We next modelled how DSB number affects the total number of crossovers for male and female meiosis in both contexts. In wild type, doubling the number of DSBs resulted in ~ 15% increase in crossovers in male and female (Figure 5G). In a context with a high number of class II crossovers, doubling the number of DSBs resulted in almost doubling the number of crossovers (Figure 5G). For a given number of DSBs the modelling predicts ~ 65% more crossovers in wt male than wt female, but essentially equal numbers of crossovers when the probability of class II crossovers is high (Figure 5G).

**Figure 5.**
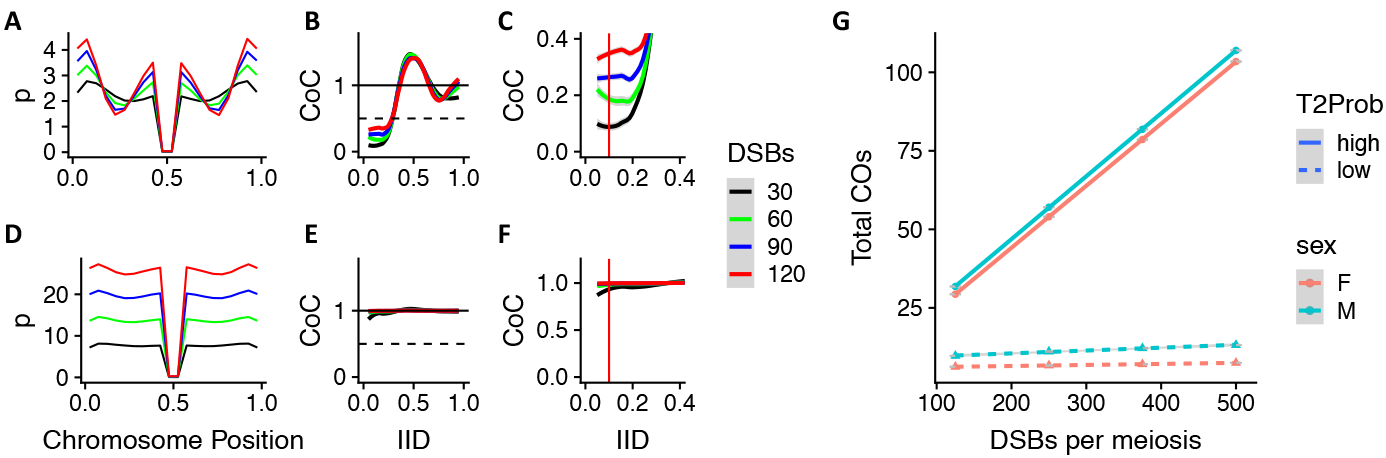
Influence of DSB number on crossover patterning. Crossover distributions (**A & D**) and CoC relationships (**B-C** and **E-F**) for beam-film simulations with varying numbers of DSBs. **A-C** show results for simulations of wt meiosis, **D-F** show results for simulation of meiosis with increased class II crossover formation (T2Prob). **(G)** Total crossover number for genome-wide simulations using best-fit parameters for male and female meiosis and varying numbers of DSBs.

## DISCUSSION

Crossover interference is a well-known genetic phenomenon, however its mechanistic basis is only just now coming to light. The interference signal is thought to propagate a set physical distance (*L*_*BF*_. usually measured in μm SC) from designated crossover sites (4, 14), and analyses commonly use cytological observations and simulations of class I crossover positions along the length of a synapsed bivalent (4, 14).

To gain insights into the differences between female and male meiosis in Arabidopsis, we analyzed a large Arabidopsis reciprocal backcross data set (16) and performed two-pathway (i.e. both class I and class II COs) beam film best-fit simulations. Our modelling suggests that the major differences in crossover number, crossover distribution and interference relationships between the sexes can all be explained by the observed difference in SC length between male and female meiosis. The relationship between genome size and SC length is governed by the size/number of chromatin loops, which occur at a conserved density of ~20 per μm SC across a wide range of organisms (28). As genome size is identical for both sexes in Arabidopsis, we would expect loop size in male meiocytes to be about 60% of that found in female meiocytes. Exactly how chromatin loop size is determined remains unclear but this decision occurs very early in, or prior to, meiosis (5, 29). It is probable, therefore, that the cause of differences in crossover patterning also occurs very early in, or prior to, meiosis. Interestingly humans also display a sex-specific differences in chromatin loop-size and SC length, although in this case female meiocytes have shorter loop-size, longer SC and more crossovers (30).

It has been reported previously that effective crossover interference is stronger in females than in males in Arabidopsis (15). Our analyses indicate that the interference signal is propagated over the same physical distance (μm SC) in both male and female meiosis, and thus from a mechanistic standpoint interference is identical in the two sexes. The higher *effective* interference (i.e. the effect on the inheritance of two linked genetic loci) observed in females can be entirely explained by the difference in SC length between the two sexes, as a given distance in μm SC corresponds to a greater length in Mb. It is worth noting that our estimates of *L*_*BF*_ for male (27.7 ± 5.6 μm) and female (23.7 ± 2.5 μm) Arabidopsis are similar to estimates for tomato (14 μm, ref 14) but are 80 to 90-fold larger than for yeast (0.3 μm, ref 14). This vast difference in the distance across which interference propagates in different taxa, as estimated by the beam-film model, remains challenging to explain biologically.

In addition to explaining differences in effective interference, SC length also explained the differences in CO distribution observed between the sexes. In male meiosis, crossovers are high adjacent to the peri-centromeres and in the distal regions, while in female meiosis crossovers are high adjacent to the peri-centromeres but low in the distal regions (3, 16). Our modelling shows that increases in the proportion of the chromosome over which interference spreads (either through a reduction in SC length, or an increase in *L*_*BF*_) reduces crossovers particularly in distal regions. The lower SC length in females can therefore account for the observed differences in crossover distribution.

In mammals, SC length is correlated with the number of DSBs (24, 25, 31). If the same holds true in plants, then we might expect fewer DSBs in female meiosis. Our analysis revealed that while the number of DSBs had very little influence on crossover distributions and CoC curves, a decrease in the number of DSBs resulted in an increase in the estimated proportion of DSB sites that become class II crossovers (*T2Prob*). Thus, the reduction in SC length observed for females, if accompanied by an equivalent reduction in DSBs, can also account for proposed differences in the number of class II crossovers between male and female meiosis. At least one line of evidence suggests this question may not be fully resolved however. In mutant lines with large numbers of additional class II crossovers, the recombination landscape of male and female meiosis are roughly equivalent with even a slightly higher number of crossovers in female (27). This suggests the possibility of similar numbers of DSBs in male and female meiosis. Further comparative cytological studies of male and female meiosis will be required to fully answer these questions.

Given the substantial differences in crossover patterning between female and male meiosis it is striking that they can all be accounted for by the difference in SC length. It is similarly striking that despite the differences in crossover patterning there are also no significant differences between the sexes in the estimated beam-film model parameters. This gives us good confidence in our approach, and suggests that similar investigations, in different contexts (e.g. mutants, over expression lines, environmental conditions), could provide further mechanistic insights into the factors governing crossover patterning in Arabidopsis.

The model can also be used to make predictions about how important agricultural goals such as heightened recombination rates could be achieved. For example, with the development of CRISPR and related technologies, it is possible to modulate the number or location of DSBs in early meiosis and there is interest in using this approach to alter recombination rates in plant breeding programs (32–34). In most organisms, crossover numbers are thought to be maintained independently from the number of DSBs through crossover homeostasis (35–37). Our modelling suggests that the extent to which homoeostasis maintains crossover numbers is determined by the proportion of DSBs that become class II crossovers: The higher the proportion of class II crossovers, the more DSB number will affect crossover number. Thus, we predict that combining the knock out of class II CO suppressing proteins (e.g. RECQ4, FANCM, FIGL1, 38–40) with approaches to increase meiotic DSBs could maximize increases in recombination and the associated benefit to breeding programs.

One of the surprising findings of our analysis is that for small inter-interval distances, an increase in the distance over which the interference signal is propagated can result in increased values of CoC (Figure 3) i.e. decreased effective interference. This behavior is not specific to the beam-film model but is expected whenever both class I and class II crossovers occur, and there is a change in the strength of suppression of closely spaced class I crossovers. This finding highlights the need for caution when interpreting interference data and particularly in the distinction between mechanistic (e.g. *L*_*BF*_) and effective (e.g. CoC from genetic data) measurements of interference. It should also be noted that at small inter-interval distances the magnitude of the predicted change in CoC is small, and that for specific interval pairs the effect of the local chromosomal landscape (e.g. recombination hotspots etc) may out-weigh the effect predicted by the model. Despite these caveats, it is clear that an increase or decrease in *mechanistic* interference strength (*L*_*BF*_) is not expected to result in an equivalent increase or decrease respectively in *effective* interference for small IIDs. Given the widespread use of reporter lines that determine recombination rates and CoC values for closely linked intervals (41) it is important to realize that these lines give little to no insight into any change in the mechanics of crossover interference.

As an example, two recent papers investigated altered recombination rates at temperature extremes in Arabidopsis (42, 43). In both cases, increased temperature gave rise to more class I COs, but the increased COs were associated with no change, or a decrease in genetic measurements of CoC (i.e. effective interference). In the studies, CoC (or interference ratio) was measured by tracking the inheritance of closely linked fluorescent reporter genes in pollen, and thus combined both class I and class II crossovers measured at a small inter-interval distance. While it could be concluded from these studies that temperature increases class I crossovers without any effect on interference, these results are also consistent with an alternative hypothesis i.e. that increased temperature decreases the distance over which interference is propagated, resulting in increased class I COs, but with no effect on genetic measurements of interference at small-inter-interval distances. Or to put it another way, high temperature might decrease *mechanistic* interference, but result in an increase (or no change) in *effective* interference for small IIDs. There is good evidence that heightened temperature might have such a mechanistic effect, given that the chromosome axis is thought to mediate interference (5) and the synaptonemal complex / axis structure is sensitive to temperature (44, 45) but this remains to be experimentally validated.

While the beam-film model was able to reliably model genetic recombination data, there are several ways in which the model might further be improved with increased understanding of the underlying biology. For example, when calculating *L*_*BF*_ and *L*_*CoC*_ in μm SC using back-cross data, we assume a direct relationship between SC length and Mb. In Arabidopsis the relationship between SC length and Mb is constant between whole chromosomes (R^2^ = 0.99, based on data from (46)), however the relationship may not be constant within a chromosome (47). Establishing how the relationship between Mb and μm SC changes for different chromosomal domains would provide one means to improve models of crossover patterning when using genetic data. Another question is whether DSB density is constant along the length of the chromosome? If so, is it constant relative to SC length or length in Mb? Recently Spo11-oligo sequencing has demonstrated relatively constant DSB formation along the length of the chromosome, although there are clearly regions of higher and lower DSB density, particularly the centromeres where DSB formation is strongly suppressed (21). Incorporating such data into the beam-film model would be an interesting future avenue.

Despite these possible improvements, it is clear that we can gain novel insights into crossover patterning using genetic recombination data in combination with beam-film simulations. These are particularly powerful when, as for this study, we have good estimates of SC length for all chromosomes, circumventing the need for cytological determination of crossover locations. This enables us to take advantage of the main benefit of genetic data, that it incorporates all crossover events, and thus enables us to develop a more nuanced understanding of the interplay between the mechanistic determinants of crossover-interference and the final effect on patterns of inheritance.

## ACKNOWLEDGEMENTS

We thank Christine Mezard for providing the experimental Arabidopsis recombination dataset and for critical reading of the manuscript. A.L. was funded by a Marie Curie fellowship (PIOF-GA-2013-628128) and the European Union’s Horizon 2020 research and innovation program under the Marie Skłodowska-Curie grant agreement No 663830. The IJPB benefits from the support of the LabEx Saclay Plant Sciences-SPS (ANR-10-LABX-0040-SPS).

## SUPPLEMENTARY TABLES AND FIGURES

**Table S1.** Chromosome metrics and beam-film parameters.

|  |  |  |  | Beam-film parameters |  |  |  |  |  |  |  |  |  |  |  |  |  |  |  |
| --- | --- | --- | --- | --- | --- | --- | --- | --- | --- | --- | --- | --- | --- | --- | --- | --- | --- | --- | --- |
| Chr | Sex | Mb | µm SC | N<sup>#</sup> | B<sup>#</sup> | E<sup>#</sup> | Bs<sup>#</sup> | Be<sup>#</sup> | Bd<sup>#</sup> | Smax<sup>^</sup> | Bsmax<sup>#</sup> | A<sup>#</sup> | L<sub>p</sub><sup>^</sup> | L<sub>mb</sub><sup>*</sup> | L<sub>sc</sub><sup>*</sup> | cL<sup>^</sup> | cR<sup>^</sup> | M<sup>#</sup> | T2prob<sup>^</sup> |
| 1 | M | 30.4 | 49.2 | 64 | 1 | 0.6 | 0.475 | 0.5 | 0.01 | 8.5 | 1 | 1 | 0.65 | 19.8 | 32.0 | 0.8 | 1 | 1 | 0.005 |
| 2 | M | 19.7 | 31.9 | 41 | 1 | 0.6 | 0.175 | 0.225 | 0.01 | 7.5 | 1 | 1 | 0.85 | 16.7 | 27.1 | 0.3 | 0.9 | 1 | 0.0065 |
| 3 | M | 23.5 | 37.9 | 49 | 1 | 0.6 | 0.5 | 0.65 | 0.01 | 5.5 | 1 | 1 | 0.7 | 16.4 | 26.6 | 0.4 | 0.9 | 1 | 0.008 |
| 4 | M | 18.6 | 30.1 | 39 | 1 | 0.6 | 0.125 | 0.225 | 0.01 | 4 | 1 | 1 | 0.6 | 11.2 | 18.0 | 0.6 | 0.9 | 1 | 0.0055 |
| 5 | M | 27.0 | 43.6 | 56 | 1 | 0.6 | 0.4 | 0.5 | 0.01 | 9.5 | 1 | 1 | 0.8 | 21.6 | 34.9 | 1.1 | 0.9 | 1 | 0.0065 |
| 1 | F | 30.4 | 25.0 | 64 | 1 | 0.6 | 0.475 | 0.5 | 0.01 | 8 | 1 | 1 | 1 | 30.4 | 25.0 | 0.7 | 0.5 | 1 | 0.003 |
| 2 | F | 19.7 | 16.2 | 41 | 1 | 0.6 | 0.175 | 0.25 | 0.01 | 7 | 1 | 1 | 1.6 | 31.5 | 25.9 | 1.2 | 0.8 | 1 | 0.004 |
| 3 | F | 23.5 | 19.3 | 49 | 1 | 0.6 | 0.5 | 0.65 | 0.01 | 6 | 1 | 1 | 1 | 23.5 | 19.3 | 0.5 | 0.6 | 1 | 0.005 |
| 4 | F | 18.6 | 15.3 | 39 | 1 | 0.6 | 0.125 | 0.225 | 0.01 | 7 | 1 | 1 | 1.7 | 31.6 | 26.0 | 0.8 | 0.5 | 1 | 0.003 |
| 5 | F | 27.0 | 22.2 | 56 | 1 | 0.6 | 0.4 | 0.5 | 0.01 | 6.5 | 1 | 1 | 1 | 27.0 | 22.2 | 0.7 | 0.6 | 1 | 0.003 |
| 1 | F<sub>DSB</sub> | 30.4 | 25.0 | 38 | 1 | 0.6 | 0.475 | 0.525 | 0.01 | 7 | 1 | 1 | 0.9 | 27.4 | 22.5 | 0.5 | 0.7 | 1 | 0.006 |
| 2 | F<sub>DSB</sub> | 19.7 | 16.2 | 38 | 1 | 0.6 | 0.475 | 0.525 | 0.01 | 6.5 | 1 | 1 | 1.5 | 29.6 | 24.3 | 1 | 0.8 | 1 | 0.004 |
| 3 | F<sub>DSB</sub> | 23.5 | 19.3 | 29 | 1 | 0.6 | 0.5 | 0.65 | 0.01 | 6 | 1 | 1 | 1 | 23.5 | 19.3 | 0.5 | 0.6 | 1 | 0.008 |
| 4 | F<sub>DSB</sub> | 18.6 | 15.3 | 23 | 1 | 0.6 | 0.125 | 0.225 | 0.01 | 6 | 1 | 1 | 1.7 | 31.6 | 26.0 | 0.8 | 0.4 | 1 | 0.005 |
| 5 | F<sub>DSB</sub> | 27.0 | 22.2 | 34 | 1 | 0.6 | 0.4 | 0.5 | 0.01 | 8.5 | 1 | 1 | 1.1 | 29.7 | 24.4 | 0.6 | 0.5 | 1 | 0.005 |

<sup>#</sup>Optimised parameter
<sup>^</sup>Fixed parameter
<sup>\*</sup>Calculated based on L<sub>p</sub>

**Figure S1.**
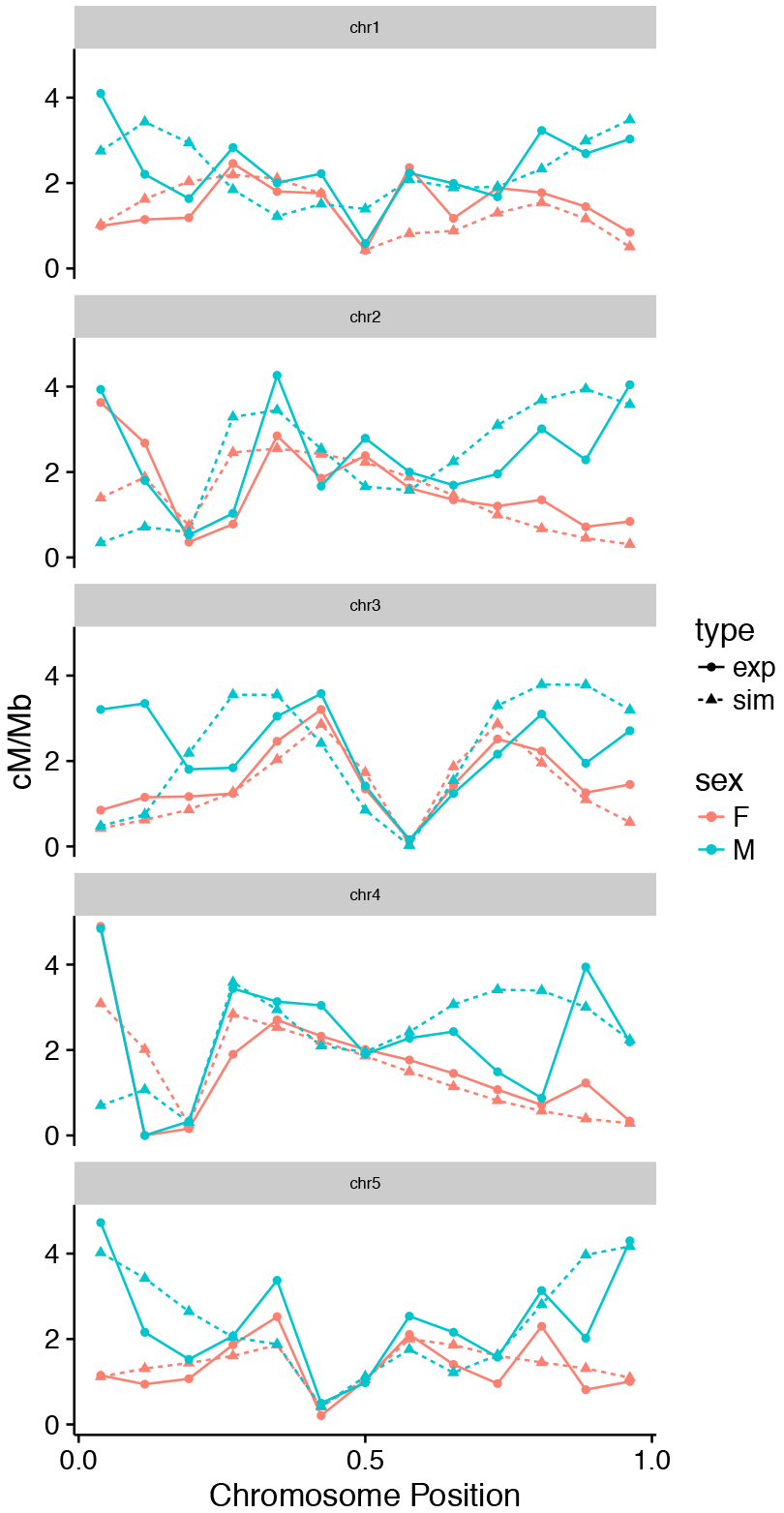
Experimental and simulated crossover distributions for Arabidopsis chromosome 1-5.

**Figure S2.**
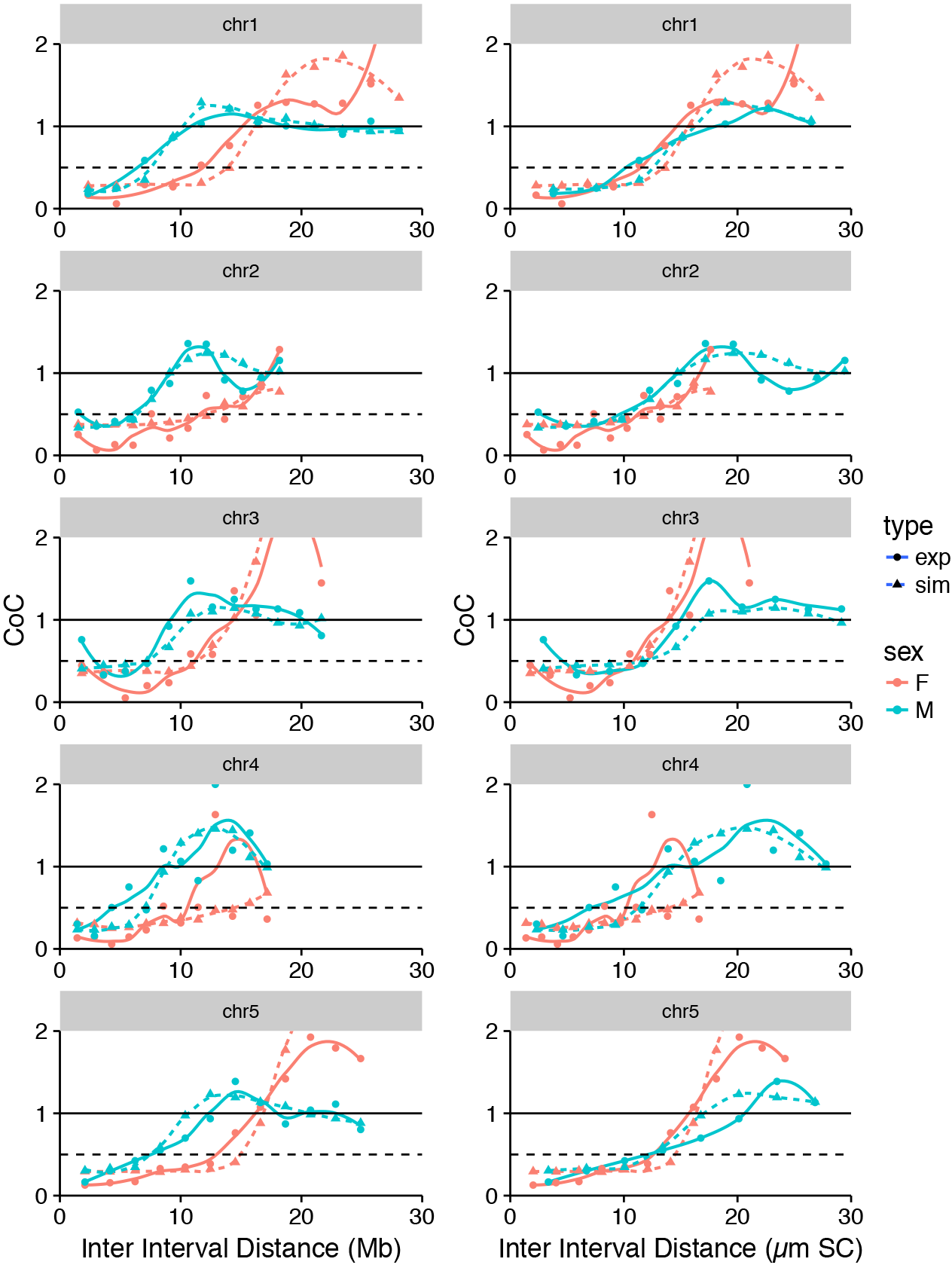
CoC curves for simulated and experimental recombination data.

**Figure 3S.**
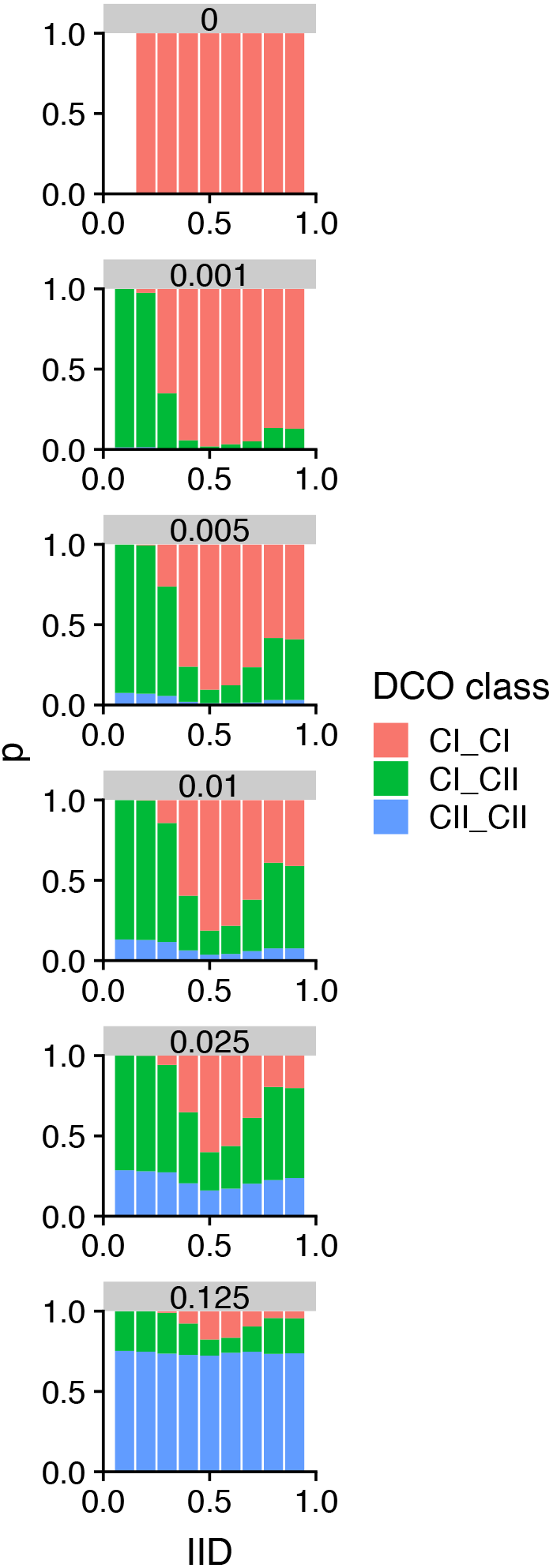
Proportions of DCOs formed between two class I crossovers (CI_CI), two class II crossovers (CII_CII), or a class I and a class II CO (CI_CII) for different IIDs and different values of T2Prob (grey bars). Total proportion of class II crossovers are as follows: T2Prob = 0, 0% class II COs; T2Prob = 0.001, 3% class II COs; T2Prob = 0.005, 13% class II COs; T2Prob = 0.01, 23% class II COs; T2Prob = 0.025, 43% class II COs; T2Prob = 0.125, 81% class II COs.

## REFERENCES

1. Sturtevant AH (1915) The behavior of the chromosomes as studied through linkage. Z Indukt Abstamm Vererbungsl 13:234–287.

2. Muller HJ (1916) The Mechanism of Crossing-Over. II. Am Nat 50:284–305.

3. Drouaud J, et al. (2007) Sex-specific crossover distributions and variations in interference level along Arabidopsis thaliana chromosome 4. PLoS Genet 3:e106.

4. Zhang L, et al. (2014) Topoisomerase II mediates meiotic crossover interference. Nature: 18–20.

5. Zickler D, Kleckner N (2015) Recombination, pairing, and synapsis of homologs during meiosis. Cold Spring Harb Perspect Biol 7:a016626.

6. Borner GV, Kleckner N, Hunter N (2004) Crossover / Noncrossover Differentiation, Synaptonemal Complex Formation, and Regulatory Surveillance at the Leptotene / Zygotene Transition of Meiosis. Cell 117:29–45.

7. Lhuissier FGP, Offenberg HH, Wittich PE, Vischer NOE, Heyting C (2007) The Mismatch Repair Protein MLH1 Marks a Subset of Strongly Interfering Crossovers in Tomato. Plant Cell Online 19:862–876.

8. Agarwal S, Roeder GS (2000) Zip3 provides a link between recombination enzymes and synaptonemal complex proteins. Cell 102:245–255.

9. Chelysheva L, et al. (2012) The Arabidopsis HEI10 is a new ZMM protein related to Zip3. PLoS Genet 8:e1002799.

10. Hollingsworth NM, Brill SJ (2004) The Mus81 solution to resolution: Generating meiotic crossovers without Holliday junctions. Genes Dev 18:117–125.

11. Kurzbauer M-T, et al. (2018) Arabidopsis thaliana FANCD2 Promotes Meiotic Crossover Formation. Plant Cell 30:tpc.00745.2017.

12. Mercier R, et al. (2005) Two Meiotic Crossover Classes Cohabit in Arabidopsis: One Is Dependent on MER3, whereas the Other One Is Not. Curr Biol 15:692–701.

13. Falque M, Anderson LK, Stack SM, Gauthier F, Martin OC (2009) Two Types of Meiotic Crossovers Coexist in Maize. Plant Cell 21:3915–3925.

14. Zhang L, Liang Z, Hutchinson J, Kleckner N (2014) Crossover patterning by the beam-film model: analysis and implications. PLoS Genet 10:e1004042.

15. Basu-roy S, Gauthier F, Giraut L, Mézard C (2013) Hot Regions of Noninterfering Crossovers Coexist with a Nonuniformly Interfering Pathway in. Genetics 195:769–779.

16. Giraut L, et al. (2011) Genome-wide crossover distribution in Arabidopsis thaliana meiosis reveals sex-specific patterns along chromosomes. PLoS Genet 7:e1002354.

17. White MA, Wang S, Zhang L, Kleckner N (2017) Quantitative Modeling and Automated Analysis of Meiotic Recombination. Meiosis, ed Stuart DT (Springer New York, New York, NY), pp 305–323.

18. White MA, Wang S, Zhang L, Kleckner N (2017) Quantitative Modeling and Automated Analysis of Meiotic Recombination. Methods Mol Biol 1471:305–323.

19. Mercier R, Mezard C, Jenczewski E, Macaisne N, Grelon M (2014) The Molecular Biology of Meiosis in Plants. Annu Rev Plant Biol 66:1–31.

20. Berchowitz LE, Copenhaver GP (2010) Genetic interference: don’t stand so close to me. Curr Genomics 11:91–102.

21. Choi K, et al. (2018) Nucleosomes and DNA methylation shape meiotic DSB frequency in Arabidopsis thaliana transposons and gene regulatory regions. Genome Res 28:1–15.

22. Kleckner N, et al. (2004) A mechanical basis for chromosome function. Proc Natl Acad Sci 101:12592–12597.

23. Panizza S, et al. (2011) Spo11-accessory proteins link double-strand break sites to the chromosome axis in early meiotic recombination. Cell 146:372–83.

24. Baier B, Hunt P, Broman KW, Hassold T (2014) Variation in Genome-Wide Levels of Meiotic Recombination Is Established at the Onset of Prophase in Mammalian Males. PLoS Genet 10. doi:10.1371/journal.pgen.1004125.

25. Gruhn JR, Rubio C, Broman KW, Hunt PA, Hassold T (2013) Cytological studies of human meiosis: Sex-specific differences in recombination originate at, or prior to, establishment of double-strand breaks. PLoS One 8:1–15.

26. Anderson LK, et al. (2014) Combined fluorescent and electron microscopic imaging unveils the specific properties of two classes of meiotic crossovers. Proc Natl Acad Sci U S A 111:13415–13420.

27. Fernandes JB, Seguéla-Arnaud M, Larchevêque C, Lloyd AH, Mercier R (2017) Unleashing meiotic crossovers in hybrid plants. Proc Natl Acad Sci:201713078.

28. Zickler D, Kleckner N (1999) Integrating Structure and Function. Annu Rev Genet 33:603–754.

29. Kaiser VB, Semple CA (2018) Chromatin loop anchors are associated with genome instability in cancer and recombination hotspots in the germline. Genome Biol 19:1–14.

30. Tease C, Hultén MA (2004) Inter-sex variation in synaptonemal complex lengths largely determine the different recombination rates in male and female germ cells. Cytogenet Genome Res 107:208–215.

31. Ruiz-Herrera A, et al. (2017) Recombination correlates with synaptonemal complex length and chromatin loop size in bovids—insights into mammalian meiotic chromosomal organization. Chromosoma 126:615–631.

32. Puchta H (2017) Applying CRISPR/Cas for genome engineering in plants: the best is yet to come. Curr Opin Plant Biol 36:1–8.

33. Hayut SF, Bessudo CM, Levy AA (2017) Targeted recombination between homologous chromosomes for precise breeding in tomato. Nat Commun 8:1–9.

34. Choi K (2017) Advances towards Controlling Meiotic Recombination for Plant Breeding. Mol Cells 40:814–822.

35. Cole F, et al. (2012) Homeostatic control of recombination is implemented progressively in mouse meiosis. Nat Cell Biol 14:424.

36. Martini E, Diaz RL, Hunter N, Keeney S (2006) Crossover Homeostasis in Yeast Meiosis. Cell 126:285–295.

37. Rosu S, Libuda DE, Villeneuve AM (2011) Robust crossover assurance and regulated interhomolog access maintain meiotic crossover number. Science 334:1286–9.

38. Crismani W, et al. (2012) FANCM Limits Meiotic Crossovers. Science 336:1588–1590.

39. Seguela-Arnaud M, et al. (2015) Multiple mechanisms limit meiotic crossovers: TOP3 α and two BLM homologs antagonize crossovers in parallel to FANCM. 112. doi:10.1073/pnas. 1423107112.

40. Girard C, et al. (2015) AAA-ATPase FIDGETIN-LIKE 1 and Helicase FANCM Antagonize Meiotic Crossovers by Distinct Mechanisms. PLOS Genet 11:e1005369.

41. Francis KE, et al. (2007) Pollen tetrad-based visual assay for meiotic recombination in Arabidopsis. Proc Natl Acad Sci U S A 104:3913–8.

42. Lloyd A, Morgan C, Franklin C, Bomblies K (2018) Plasticity of meiotic recombination rates in response to temperature in Arabidopsis.

43. Modliszewski JL, et al. (2018) Elevated temperature increases meiotic crossover frequency via the interfering (Type I) pathway in Arabidopsis thaliana. PLoS Genet 14:1–15.

44. Loidl J (1989) Effects of elevated temperature on meiotic chromosome synapsis in Allium ursinum. Chromosoma 97:449–458.

45. Morgan CH, Zhang H, Bomblies K (2017) Are the effects of elevated temperature on meiotic recombination and thermotolerance linked via the axis and synaptonemal complex? Philos Trans R Soc B Biol Sci 372. doi:10.1098/rstb.2016.0470.

46. López E, Pradillo M, Romero C, Santos JL, Cuñado N (2008) Pairing and synapsis in wild type Arabidopsis thaliana. Chromosom Res 16:701–708.

47. Bickmore WA, Oghene K (1996) Visualizing the spatial relationships between defined DNA sequences and the axial region of extracted metaphase chromosomes. Cell 84:95–104.

